# Multi-omic Evaluations Nominate an ER-Mitochondrial Axis and Inflammatory Macrophage as Drivers of Right Atrial Dysfunction

**DOI:** 10.1101/2025.03.22.644722

**Authors:** Jenna B. Mendelson, Jacob D. Sternbach, Jeffrey C. Blake, Minwoo Kim, Ryan A. Moon, Rashmi M. Raveendran, Lynn M. Hartweck, Walt Tollison, Matthew Lahiri, John P. Carney, Todd Markowski, LeeAnn Higgins, Cutler T. Lewandowski, Felipe Kazmirczak, Gaurav Choudhary, Kurt. W. Prins

## Abstract

**Background:** Right atrial (RA) dysfunction is an emerging risk factor for poor outcomes in pulmonary arterial hypertension, however the mechanisms underlying compromised RA function are understudied.

**Objectives:** Multi-omic analyses defined the cellular and molecular mediators associated with RA dysfunction in pulmonary artery banded (PAB) swine.

**Methods:** 4-week-old castrated male Yorkshire pigs were subjected to PAB and aged six weeks to induce right heart failure. Cardiac MRI evaluated RA size and function. snRNAseq defined the cell-specific alterations in RA tissue. Mitochondrial proteomics and metabolomics analyses examined the metabolic alterations in RA samples. Inducible pluripotent stem cell-derived atrial cardiomyocytes (iPSC-ACM) were treated with tunicamycin to induce endoplasmic reticulum (ER) stress and mitochondrial structure and function were probed.

**Results:** PAB induced RA dilation/dysfunction and atrial cardiomyocyte hypertrophy. snRNAseq demonstrated PAB altered the cellular composition of the RA defined by increased inflammatory macrophages and an alteration of cardiomyocyte subpopulations. RA cardiomyocytes exhibited ER stress and mitochondrial metabolic enzyme dysregulation. PAB RAs, but not PAB right ventricles, had downregulation of branched chain amino acid degrading enzymes. Metabolomics profiling revealed BCAA and fatty acid metabolism were impaired in the dysfunctional RA. Tunicamycin-induced ER stress disrupted mitochondrial structure/function in iPSC-ACMs.

**Conclusions:** Multi-omic evaluations demonstrate RA dysfunction is characterized by cardiomyocyte metabolic derangements due to ER dysregulation and an accumulation of pro-inflammatory macrophages.

## Introduction

Right heart failure is a leading cause of morbidity and mortality in numerous cardiovascular diseases(1,2). Right heart function is reliant on the synergistic actions of the right atrium (RA) and the right ventricle (RV) to facilitate the circulation of deoxygenated venous blood through the pulmonary vasculature and then fill the left ventricle(3). Recent studies demonstrate RA dysfunction is an emerging risk factor for heightened mortality in pulmonary arterial hypertension (PAH)(4,5), a disease where right heart failure is the leading cause of death(1). Moreover, the RA appears to be a sensitive marker of compromised right heart function as RA dilation occurs before overt RV dilation or dysfunction are detectable in multiple etiologies of pulmonary hypertension(6). While available data suggest the RA and RV work in concert to optimize right heart function, the mechanisms underlying RA dysfunction are vastly understudied as compared to our growing understanding of RV failure(3).

Recent data from human studies are providing new physiological and molecular insights into the consequences of RA dysfunction in PAH. First, RV diastolic stiffness compromises RA function, leading to pathological vena cava backflow(7). Second, PAH patients have RA cardiomyocyte hypertrophy and heightened atrial fibrosis, and these changes are associated with impaired RA-RV coupling(8). In addition to suppressed pump function, the compromised RA negatively impacts right heart function via atrial arrhythmias(9). The presence of atrial arrhythmias is prognostically important as they are associated with greater morbidity and mortality in PAH(9). Certainly, these data highlight the importance of the RA in patient outcomes and alterations that occur with RA dysfunction, but an unbiased and deep molecular characterization of dysfunctional RA tissue is lacking.

Here, we addressed these key knowledge gaps by delineating how pulmonary artery banding modulated RA structure and function in pigs. Then, we performed a multi-omic evaluation of RA tissue that included single nucleus RNAseq (snRNAseq), proteomics, and metabolomics to define the cellular and molecular alterations that occurred in the dysfunctional RA.

## Methods

A detailed description of all methods is provided in the Supplemental Methods.

Experimental animals and physiological analysis: Control and PAB animals were previously described(10). Cardiac MRI and hemodynamic evaluations were conducted as previously discussed(10).

Multi-omic evaluation: We performed single nucleus RNAsequencing(10), mitochondrial proteomics(10,11), and metabolomics analysis in collaboration with Metabolon to define the molecular signature of the dysfunctional right atrium.

iPSC-ACM culture and mitochondrial assessments: Human inducible pluripotent cells (Allen Institute) were differentiated into atrial cardiomyocytes using the atrial cardiomyocyte differentiation kit (StemCell Technologies). After differentiation, iPSC-ACM were treated with sham or tunicamycin, then cells were stained with MitoTracker Orange (ThermoScientific) and network morphology was examined using a Zeiss LSM900 Airyscan 2.0 microscope. Mitochondrial function of iPSC-ACM was evaluated on a Seahorse XF96(12).

Statistical Analysis: Results are presented as mean±standard deviation if normally distributed or median (25^th^, 75^th^ percentiles) if not normally distributed as evaluated by Shapiro-Wilk test. Differences between two groups were determined by unpaired t-test (normal distribution and equivalent variance) or Mann-Whitney test. A p-value <0.05 was considered significant. Statistical analyses were performed on GraphPad version 10.0.

## Results

### Pulmonary artery banded pigs exhibited RA dilation, hypocontractility, and cardiomyocyte hypertrophy

Cardiac MRI defined RA size and function in five control and five PAB pigs (**Figure 1A**). PAB-pigs exhibited significant RA remodeling with increased RA end-diastolic volume (**Figure 1B**) and reduced atrial function as RA ejection fraction and the right atrioventricular coupling index was significantly altered (**Figure 1 C and D**). Next, we evaluated the histological response of atrial cardiomyocytes to PAB and found RA cardiomyocyte hypertrophy as there was a >two-fold increase in cardiomyocyte cross sectional area in PAB animals (**Figure 1 E and F**). These data demonstrated PAB resulted in RA remodeling and dysfunction.

**Figure 1:**
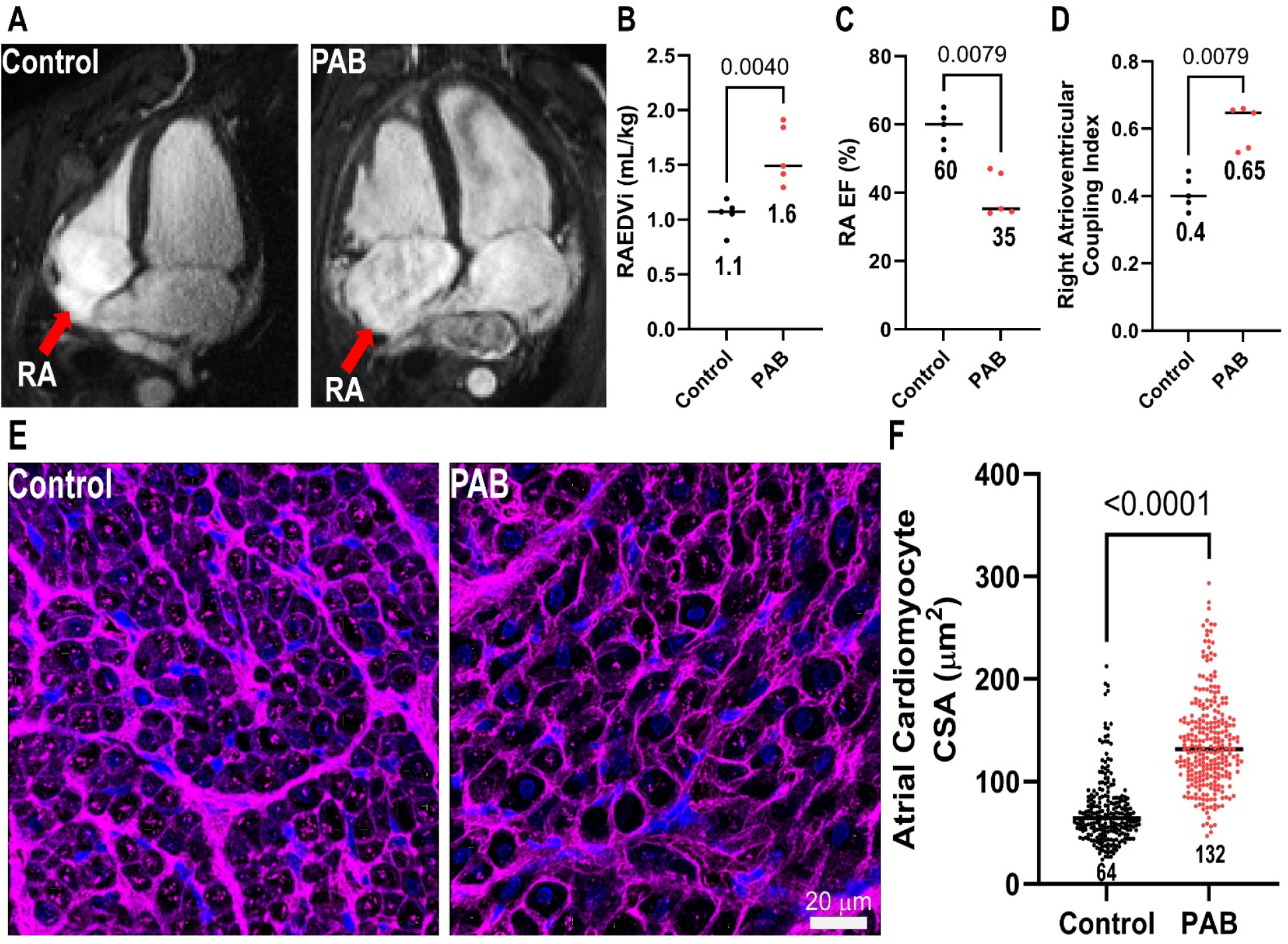
Pulmonary artery banding induced RA dilation, dysfunction, and RA cardiomyocyte hypertrophy. (A) Representative 4-chamber cMRI images depicting RA enlargement in PAB pigs. (B) Quantification of RA end-diastolic volume index (Control: 1.1±0.1 mL/kg, PAB: 1.6±0.3 mL/kg). *p*-value determined by unpaired *t*-test. (C) Quantification of RA ejection fraction [Control: 60 (54, 63), PAB: 35 (34, 46)] *p*-value determined by Mann-Whitney test. (D) Quantification of right atrioventricular coupling index [Control: 0.40 (0.37, 0.47), PAB: 0.65 (0.54, 0.66)] *p*-value determined by Mann-Whitney test (E) Representative confocal micrographs of control and PAB RA cross sections stained with wheat germ agglutinin (purple) and 4’,6-diamidino-2-phenylindole to delineate nuclei (blue). (E) Quantification of RA cardiomyocyte cross sectional area [Control: 64 *µ*m^2^ (51, 84), PAB: 132 *µ*m^2^ (106, 160)], *p*-value determined by Mann-Whitney test.

### snRNA-seq defined the cellular landscape of the right atrium and delineated cell-specific alterations

We used snRNAseq to evaluate the cellular composition and define cell-specific alterations in RA specimens (**Figure 2 A and B**). After filtering data, we compared 55,152 nuclei from four control RAs and 63,878 nuclei from four PAB RAs. We identified eight cell types (lymphocytes, fibroblasts, macrophages, endothelial cells, cardiomyocytes, adipocytes, neuronal cells, and pericytes) in RA samples (**Figure 2 C and D**). When comparing the cellular composition between the two groups, PAB RA samples had increased relative abundances of macrophage and a reduction in fibroblast abundances (**Figure 2 C and D**). Then, we determined cell-specific alterations in transcriptional regulation. The cells with the highest number of differentially expressed genes (DEG) were cardiomyocytes followed by macrophages. Adipocytes, neural cells, and pericytes did not have any DEGs (**Figure 2E**).

**Figure 2:**
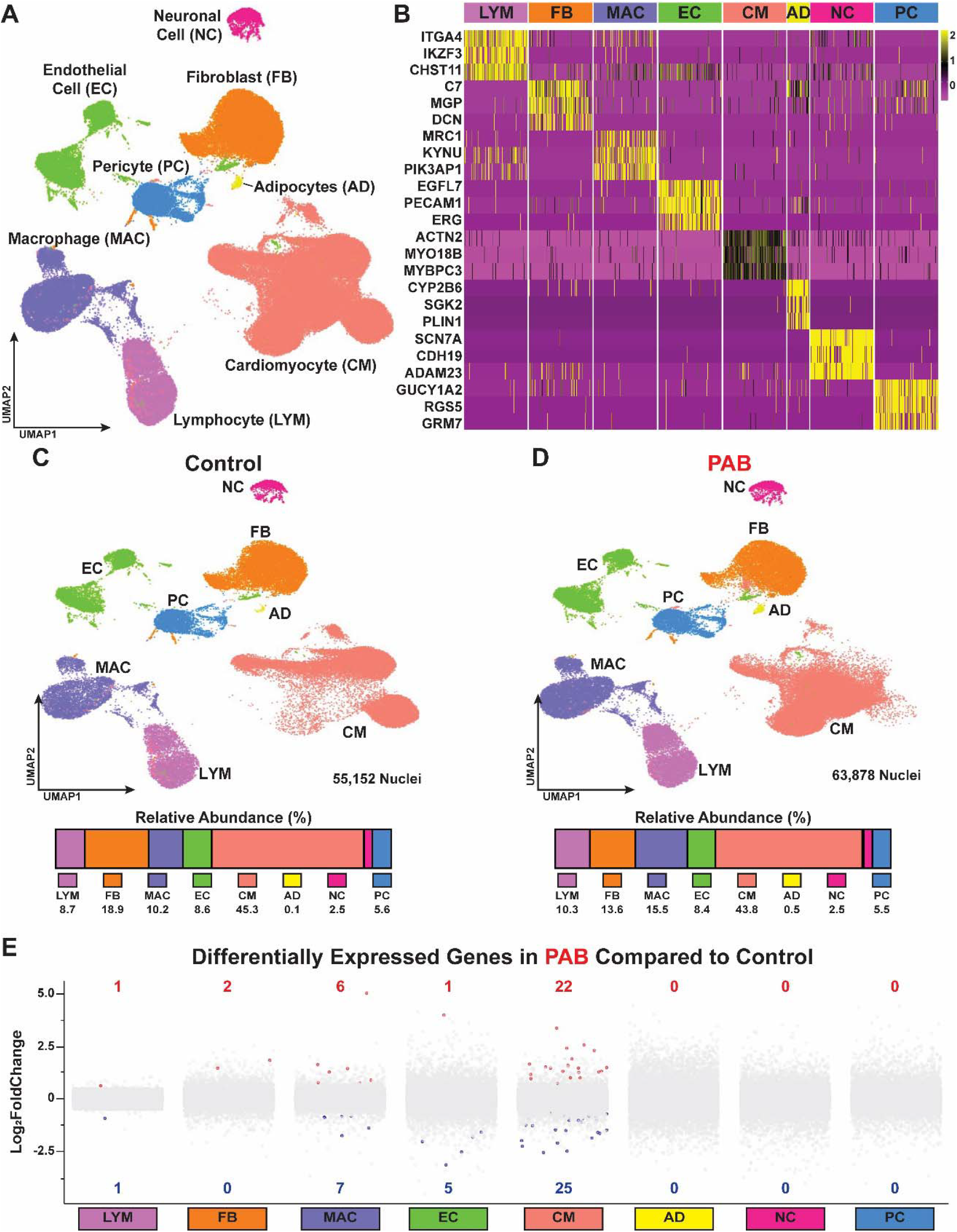
snRNAseq delineated the cell-specific alterations in the dysfunctional right atrium. (A) UMAP depicting the eight cluster of cells detected in RA samples. (B) Heatmap demonstrating unique transcriptomic profile of each cell cluster. (C and D) UMAP of control and PAB samples with proportion of each cell type identified in each experimental group. (E) Depiction of the differentially expressed genes in each cell type.

### Cardiomyocyte derangements were present in the dysfunctional right atrium

We then probed the cardiomyocyte (CM) population of the RA to better understand why the PAB RA was hypocontractile. Unbiased clustering identified eight CM subtypes in the RA, and there were distinct differences in the cardiomyocyte composition as PAB animals had lower abundances of CM2 and higher abundances of CM1 and CM3 (**Figure 3 A-D**). Pathway analysis of the upregulated transcripts identified alterations in the metabolism of amino acids and bile acids (**Figure 3E**). When evaluating downregulated transcripts, the only pathway enriched was the endoplasmic reticulum (ER) stress pathway (**Figure 3F**).

**Figure 3:**
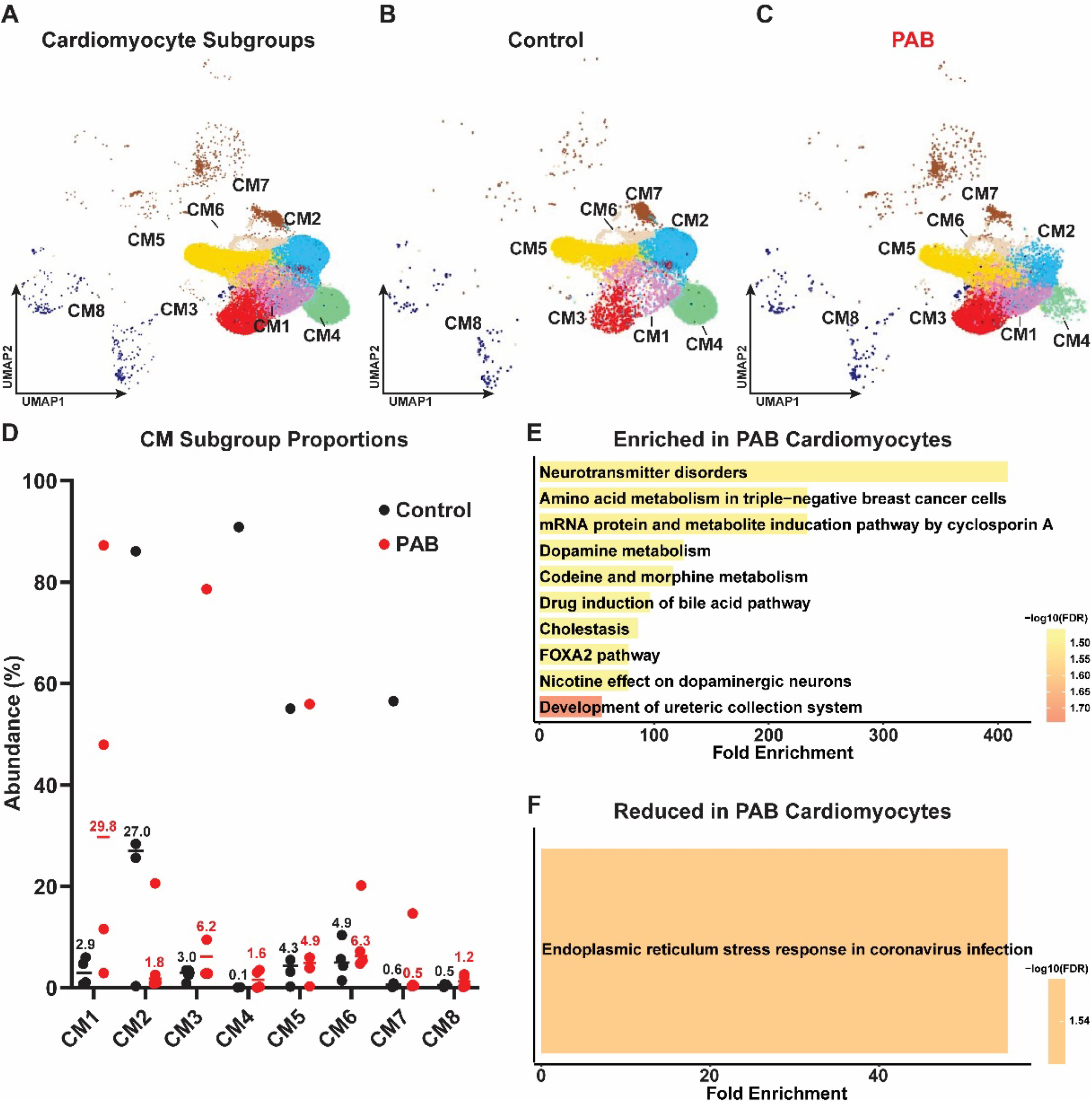
PAB shifted and cardiomyocyte subcluster populations and induced disruptions in metabolism and endoplasmic reticulum homeostasis in RA cardiomyocytes. (A) UMAP of combined cardiomyocyte clusters. (B) UMAP of control cardiomyocyte cells. (D) UMAP of PAB cardiomyocyte clusters. (D) Quantification of relative abundances of each cardiomyocyte subpopulation. (E) Pathway analysis of transcripts that were significantly elevated in PAB cardiomyocytes. (F) Pathway analysis of downregulated transcripts in PAB cardiomyocytes.

To further examine if ER homeostasis was altered in the dysfunctional RA, we probed numerous ER protein abundances with immunoblots. We observed downregulation of all nearly ER proteins in PAB RAs (**Supplemental Figure 1**). These data, plus our RNAseq analysis, provided strong evidence that ER homeostasis was impaired in the PAB RA.

### PAB increased the abundance of pro-inflammatory macrophage in the RA

Because the macrophage relative abundance change was the numerically greatest of all cell types in our snRNAseq analysis and they had the second most DEGs, we further delineated how PAB modulated macrophage subtypes and molecular regulation. Both control and PAB RAs had three distinct macrophage populations, and the relative abundances of each macrophage subpopulation were equivalent (**Supplemental Figure 2A-C**). However, PAB RA samples had greater numbers of each macrophage in all subclusters (**Supplemental Figure 2D**). When performing pathway analysis of DEGs, we found the PAB macrophage exhibited a pro-inflammatory phenotype as numerous inflammatory pathways were enriched in the upregulated genes while metabolic pathways implicated in macrophage inflammatory response(13) (one-carbon metabolism and folic acid) were downregulated (**Supplemental Figure 2E-F**). Cell chat analysis predicted the number of interactions between PAB macrophage and other cell types were reduced, but the strength of interactions between macrophage and cardiomyocytes was increased (**Supplemental Figure 2G-H**).

### Proteomics identified impaired branched chain amino acid (BCAA) catabolism in the RA but not the RV of PAB pigs

We next performed a proteomics analysis of mitochondrial enrichments in the RA and RV of four control and four PAB animals to determine if there were RA-specific pathways associated with impaired chamber function. Hierarchical cluster analysis revealed each tissue had a distinct proteomic signature and there were several differences between control and PAB RA samples (**Figure 4A**). To define which proteins were associated with specific chamber function, we performed correlational analysis to delineate which pathways were associated with RA or RV dysfunction. Pathway analysis of proteins enriched with greater RA or RV ejection fraction were mainly metabolic, and there was significant overlap between the two chambers. Interestingly, proteins responsible for BCAA degradation were uniquely associated with RA but not RV ejection fraction (**Figure 4B**). Protein pathways associated with chamber dysfunction were mostly similar and included ribosomes, glycolysis/gluconeogenesis, and hypoxia-inducible factor 1a (HIF1) signaling (**Figure 4C**), but there were more inflammatory pathways identified in the RA than the RV.

**Figure 4:**
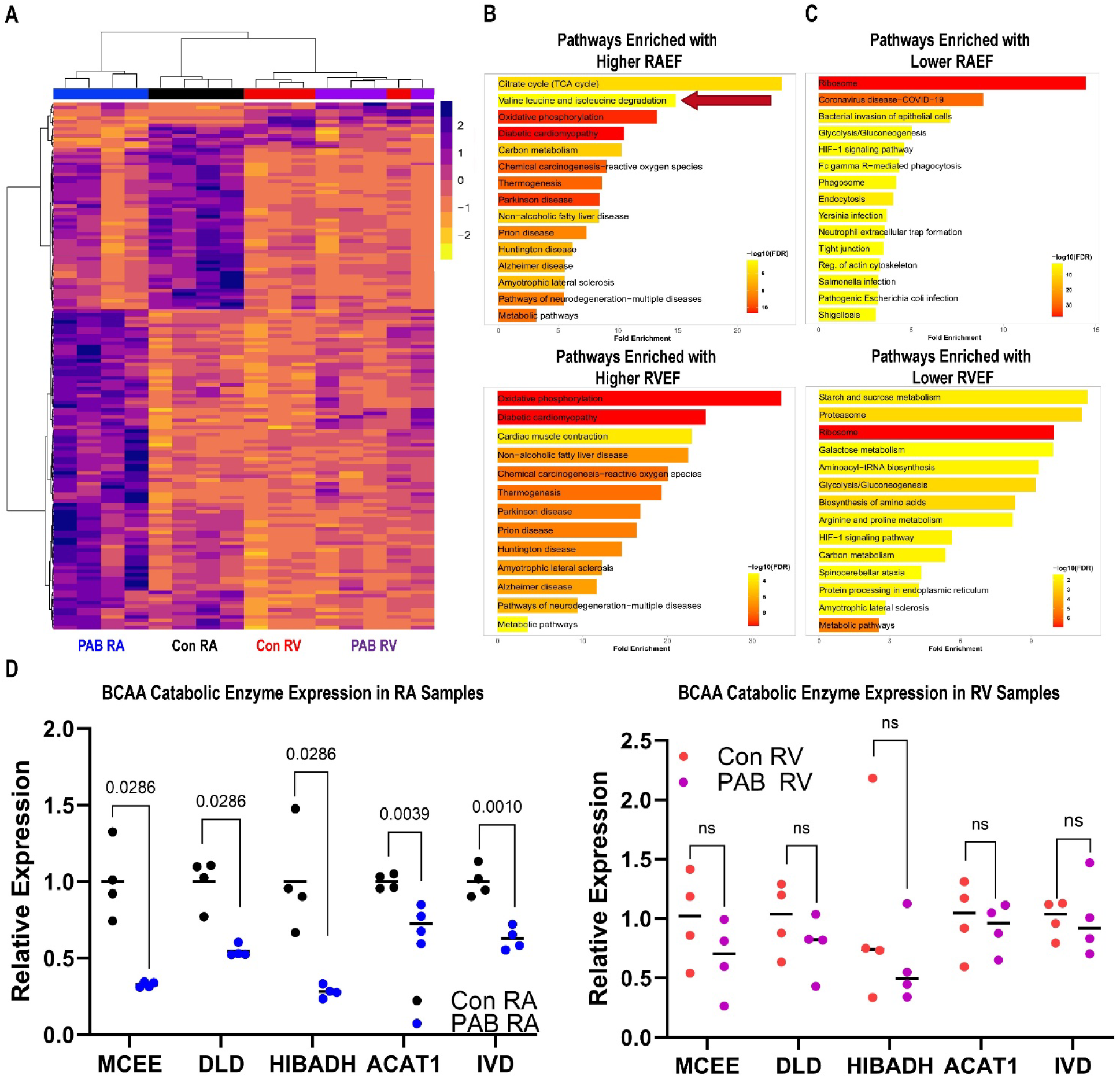
Proteomics profiling defined disruptions in BCAA catabolism in an atrial-specific manner. (A) Hierarchical cluster analysis of top 150 differentially expressed proteins in mitochondrial enrichments from RA and RV samples. (B) Pathway analysis of proteins associated with higher RA ejection fraction (top) and higher RV ejection fraction (bottom). Proteins associated with BCAA catabolism were uniquely regulated in the RA. (C) Pathway analysis of proteins associated with impaired RA ejection fraction (top) and RV ejection fraction (bottom). (D) Quantification of five BCAA catabolic proteins from RA and RV samples. BCAA catabolic proteins were uniquely downregulated in the RA but not the RV. *p*-values determined by Mann-Whitney or unpaired t-tests.

To further probe the BCAA pathway in a chamber-specific fashion, we evaluated how the five BCAA catabolism enzymes [methylmalonyl-CoA epimerase (MCEE), dihydrolipoamide dehydrogenase (DLD), 3-hydroxyisobutyrate dehydrogenase (HIBADH), acetyl-CoA acetyltransferase 1 (ACAT1), isovaleryl-CoA dehydrogenase (IVD)] significantly associated with RA ejection fraction were regulated in the RA and RV. All five enzymes were significantly reduced in PAB RA samples, but none were reduced in PAB RV samples (**Figure 4D**). Thus, these data highlighted metabolic alterations were associated with RA and RV dysfunction, but nominated BCAA metabolism as a RA-specific finding.

### Metabolomics profiling identified alterations in BCAA and fatty acid metabolites in the PAB RA

Next, we used global metabolomics profiling to probe the functional consequences of mitochondrial metabolic protein dysregulation in the RA. Hierarchical cluster analysis demonstrated the two experimental groups had relatively distinct metabolomics profiles (**Figure 5A**). Next, we performed correlational analysis to define what metabolites and their subsequent pathways were associated with either higher or lower RA ejection fraction. Metabolite pathways enriched with higher RA ejection fraction included amino acid (histidine, alanine, aspartate, and glutamate) metabolism, TCA cycle, glutathione metabolism, and starch/sucrose metabolism (**Figure 5B**). Pathways enriched with lower RA ejection fraction included aromatic amino acids, BCAA, and fatty acid metabolism (**Figure 5C**). Then, we specifically profiled BCAA-associated metabolites because both our proteomics and metabolomics analyses suggested this pathway was dysregulated in PAB RAs. Nearly all BCAA-associated metabolites were elevated in PAB RA samples (**Figure 5D**). Furthermore, we evaluated relative abundances of fatty acid metabolites based on findings from our proteomics and metabolomics pathway analyses. Both long chain fatty acids and acylcarnitines accumulated in PAB RA samples (**Figure 5 E and F**), which suggested diseased RA cardiomyocytes were unable to fully oxidize fatty acids.

**Figure 5:**
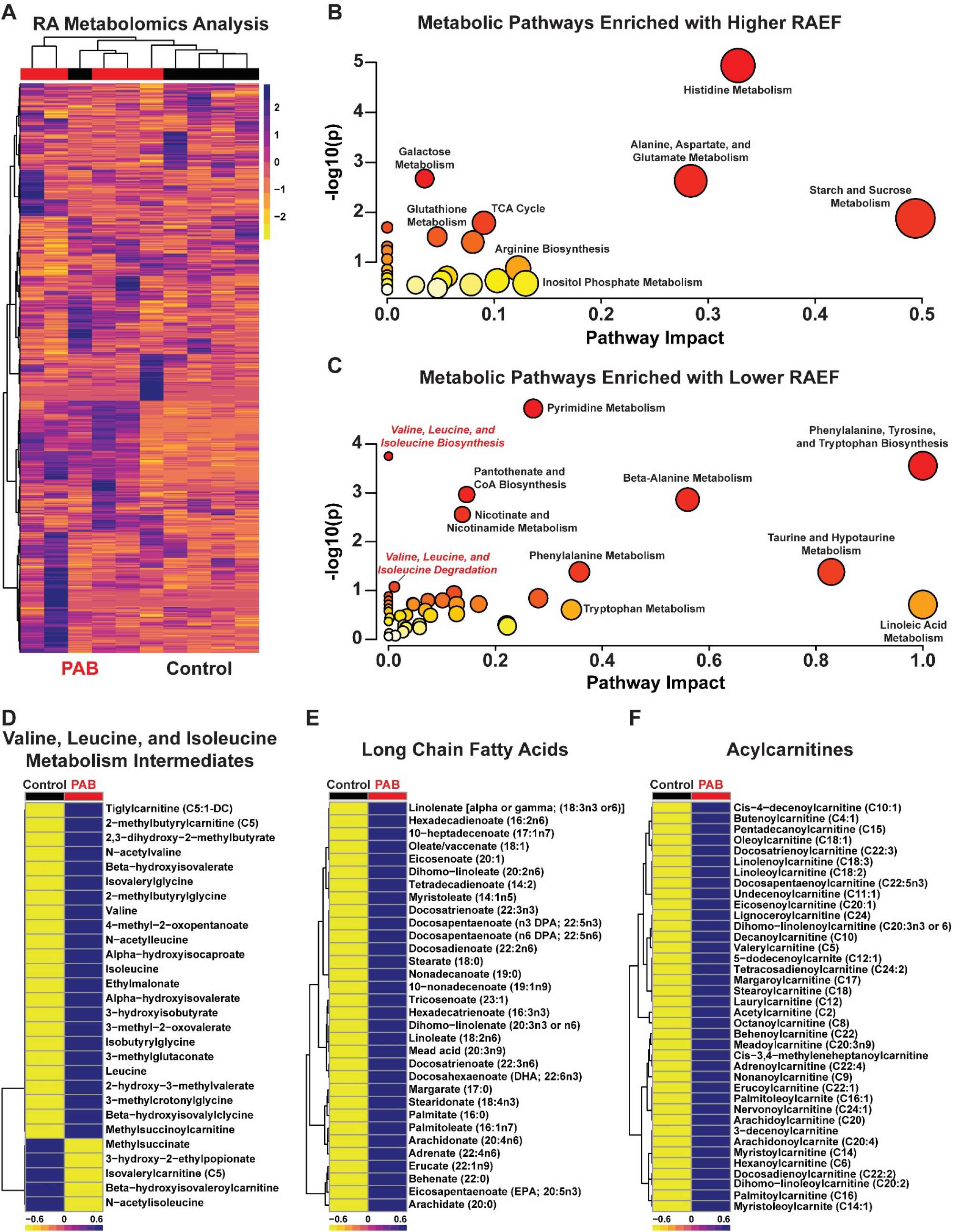
Metabolomics analysis defined alterations in BCAA and fatty acid metabolism in the dysfunctional RA. (A) Hierarchical cluster analysis of 690 metabolites in five control and five PAB RA samples. (B) Depiction of metabolic pathways positively associated with RA ejection fraction. (C) Depiction of metabolic pathways inversely associated with RA ejection fraction. Pathways included BCAA metabolism and fatty acid metabolism. (D) Hierarchical cluster analysis revealed an accumulation of nearly all BCAA associated metabolites in PAB RA samples. Impairments in fatty acid metabolism were suggested as long chain fatty acids (E) and acylcarnitines (F) accumulated in PAB RA samples.

### Electron microscopy defined alterations in RA cardiomyocyte mitochondria

To expand on our proteomics and metabolomics analyses, we analyzed electron micrographs of mitochondria in RA cardiomyocytes to determine how mitochondrial morphology was impacted by PAB. There was no significant difference in mitochondrial length:width ratio in PAB mitochondria, but mitochondrial cristae morphology was compromised in PAB mitochondria (**Supplemental Figure 3A-B**). Thus, RA cardiomyocytes exhibited mitochondrial morphological changes, proteomic dysregulation, and metabolomic alterations in PAB RAs.

#### Induction of ER stress compromised mitochondrial function in iPSC-ACM

Finally, we evaluated how disruption of ER homeostasis impacted mitochondrial structure and function to nominate a potentially targetable pathway that integrates our multi-omics analyses. To do so, we examined how treatment of iPSC-ACMs (**Supplemental Figure 4A**) with tunicamycin, an ER poison (14,15), impacted mitochondrial structure and function. Tunicamycin disrupted mitochondrial network morphology (**Supplemental Figure 4 B and C**). Seahorse analysis demonstrated tunicamycin impaired iPSC-ACM mitochondrial metabolism as baseline, maximal, and ATP-linked oxygen consumption rates were significantly reduced (**Supplemental Figure 4 D-F**). In conclusion, these data demonstrated impairments in ER homeostasis disrupted mitochondrial structure and function in iPSC-ACMs, which aligned with our multi-omic evaluations.

## Discussion

Here, we provide the first-ever multi-omic evaluation of the dysfunctional RA in the porcine PAB model of right heart failure. We demonstrate PAB induces RA dilation and dysfunction using cardiac MRI analysis. Histologically, PAB atrial cardiomyocytes are hypertrophied with a >two-fold increase in cross sectional area. Using snRNAseq, we delineate cell-type specific alterations that occur in RA myopathy, and when specifically profiling cardiomyocytes we find ER homeostasis is compromised. In addition, we show macrophage relative abundances increase in the PAB RA, and pathway analysis suggests these macrophages are pro-inflammatory. Proteomics profiling identifies reductions in electron transport chain and fatty acid oxidation proteins in the dysfunctional RA, and we demonstrate an atrial-specific dysregulation of BCAA catabolic enzymes. Metabolomics analysis corroborates our proteomics data as PAB RA samples accumulate BCAA metabolites, long chain fatty acids, and acylcarnitines. Finally, induction of ER stress in iPSC-ACMs alters mitochondrial structure and function. The summation of our data suggests both an ER-mitochondrial atrial cardiomyocyte metabolic axis and an accrual of pro-inflammatory macrophage as mechanistic drivers of RA dysfunction (**Figure 6**).

**Figure 6:**
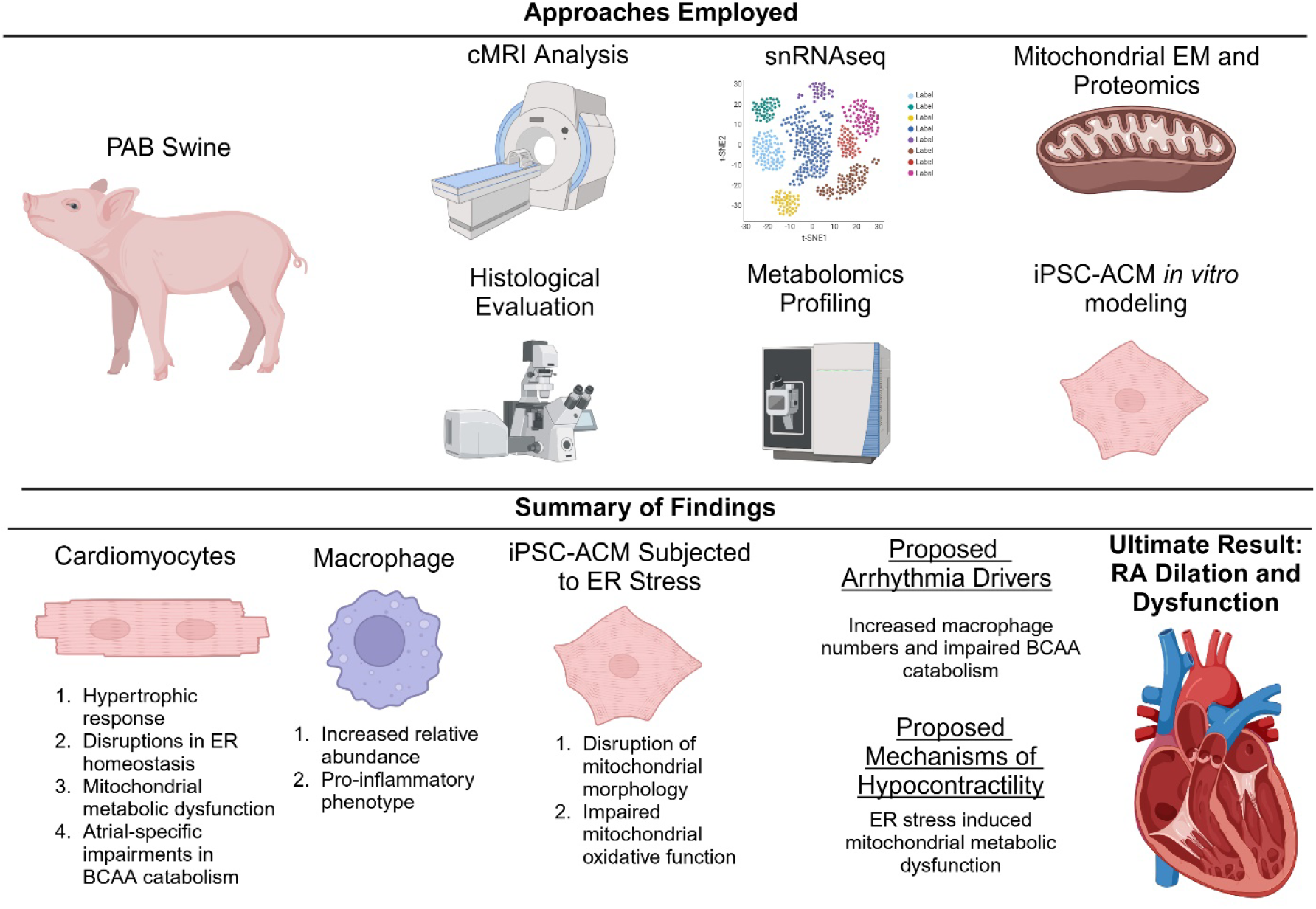
Central Figure: Approaches implemented, summary of findings, and proposed cellular and molecular mechanisms driving RA dysfunction.

Although relative abundances of cardiomyocytes are not reduced in PAB RA, which is distinct from what we observe in the dysfunctional porcine RV(10), there are significant alterations in cardiomyocyte composition and the metabolic profile of RA cardiomyocytes. In particular, PAB RAs have downregulation of BCAA degrading enzymes, which is accompanied with an accumulation of BCAA metabolites. Moreover, there is a positive association between RA ejection fraction and oxidative phosphorylation and fatty acid oxidation enzymes. These enzymatic changes may explain why long chain fatty acids and acylcarnitines accumulate in the PAB RA as fatty acid metabolism relies both on fatty acid oxidation enzymes and the electron transport chain(16). Perhaps, approaches that enhance mitochondrial biogenesis and normalize BCAA catabolism, oxidative phosphorylation proteins, and fatty acid oxidation enzymes would counteract RA hypocontractility.

The cell type with the greatest change in abundance and the second most differentially expressed genes in the diseased RA are macrophages. Interestingly, macrophage relative abundance is higher in RA samples than what we previously found in RV samples(10), which is congruent with data from the human heart cell atlas(17), and implies macrophages may have pronounced effects on atrial function. Following PAB, atrial macrophage relative abundance increases, and they possess a pro-inflammatory molecular phenotype, suggesting they may contribute to the observed atrial myopathy. An accumulation of pro-inflammatory macrophages occurs in the left atrium in patients and rodents with atrial fibrillation(18,19). Suppressing the pro-inflammatory phenotype of macrophages mitigates atrial fibrillation in rodents, which provides evidence that macrophage modulate atrial function(18). Further evaluations of macrophage targeting strategies should be performed to determine if this would be a novel approach to correct RA function.

Our data suggest the ER stress pathway is impaired in the dysfunctional RA, which may serve as an important therapeutic target to augment RA function. Previous studies demonstrate ER stress promotes atrial fibrillation as the ER stress pathway is dysregulated in rodents and humans with atrial fibrillation. Moreover, small molecule antagonism of the ER stress pathway reduces atrial fibrillation induction in rodents(20,21). Although these studies demonstrate counteracting ER stress improves left atrium physiology, it is possible that these findings would hold true in the RA. Interestingly, global induction of ER stress pathway results in a cardiomyopathy defined by mitochondrial morphological deficits and metabolic derangements(22). Unfortunately, atrial structure and function were not examined in this study, so it is presently unclear if modulating ER stress is sufficient to disrupt atrial physiology. Nonetheless, these data fit well with our Seahorse data showing induction of ER stress disrupts mitochondrial structure and function in iPSC-ACM (**Supplemental Figure 4**), and ultimately imply ER dysfunction promotes mitochondrial metabolic derangements in the diseased RA.

We nominate BCAA catabolism as a RA-specific finding as these enzymes are only downregulated in RA samples, and this finding may have therapeutic implications for right heart function. Although not evaluated in the setting of RA dysfunction, elevated BCAAs are observed in the atrial appendage of atrial fibrillation patients(23). Furthermore, excess BCAAs induce pro-arrhythmic effects in human iPSC-CMs(24). Our observation of an atrial-specific dysregulation of BCAA catabolic enzymes may shed more light into the controversial field of BCAAs in heart failure. While small molecule augmentation of BCAA catabolism improves cardiac function in aortic banded rodents(25), cardiac muscle specific enhancement of BCAA metabolism has minimal effects on cardiac physiology following aortic banding(26). Murashige *et al*., suggest enhancing BCAA catabolism exerts cardioprotective effects via a reduction in systemic vascular resistance. However, neither of these two manuscripts examined how augmenting BCAA metabolism impacted atrial function in their studies. Perhaps some of the cardiovascular benefits of augmenting BCAA catabolism is from improved atrial function.

While it is possible that the atria more readily metabolize BCAAs for ATP generation, analysis of human cell atlas and our proteomics data do not suggest BCAA catabolic enzymes are higher in the RA than the RV (**Supplemental Figure 5**), but substrate delivery could also contribute to differences in BCAA metabolism between the two chambers. Interestingly, there is precedence that ER stress alters BCAA metabolism as tunicamycin treatment downregulates BCAA enzymes in adipocytes(27). Thus, ER stress may underlie the BCAA phenotype we observe in diseased RA cardiomyocytes.

Finally, there is emerging data suggesting the RA modulates systemic biology in PAH via its ability to synthesize and secrete bone morphogenic protein 10 (BMP10), a signaling molecule that regulates endothelial cell function via the transforming growth factor-β pathway(28). In human PAH samples, BMP10 mRNA and protein levels are increased in the RA(29). In our porcine samples, we find BMP10 mRNA is almost exclusively detectable in cardiomyocytes, however BMP10 expression is not significantly elevated in PAB RA cardiomyocytes (**Supplemental Figure 6**). This could reflect sex or species differences as the previous human study included both females and males. Perhaps future therapeutic studies counteracting RA dysfunction should be probed for BMP10 expression because BMP10 is an important biomarker for RA structure/function and it could modulate systemic endothelial cell biology.

### Limitations

Our study has important limitations that we acknowledge. First, this study was performed in castrated young males, and thus it does not reflect adult disease and lacks an evaluation of females. Second, we did not perform an intervention so we do not know if any of our identified pathways can be targeted to improve RA function. Third, we did not see significant alterations in fibroblasts but previous studies show RA fibrosis is increased in PAH RA(8). This may be explained by our use of castrated animals because castration can mitigate fibrosis in the RV of rodents(30). Unfortunately, we were unable to detect multiple phosphorylated ER stress proteins on Western blot, which prevented our ability to evaluate canonical ER stress proteins (phospho-XBP and phosphor-IRE). Finally, we did not detect any atrial arrhythmias in our porcine model, but they were only evaluated during hemodynamic assessments and cardiac MRI at the end of the study. Future studies examining atrial arrhythmia inducibility are needed.

