## Supplemental Data for "Multi-omic Evaluations Nominate an ER-Mitochondrial Axis and Inflammatory Macrophage as Drivers of Right Atrial Dysfunction"

**Supplemental Methods**

Animals: Pulmonary artery banding (PAB) was performed as previously described(1-4) on 5 Yorkshire castrated male pigs aged 28-35 days old and weighing 6.5±1.4 kg on the day of surgery. Five castrated pigs of a similar age and weight were used as controls, and were housed at the University of Minnesota Research Animal Resources Facility for six weeks without surgical intervention.

Cardiac MRI (cMRI) Examination: cMRI studies were performed on a Siemens Aera 1.5T scanner as previously described, and analyzed by an experienced cardiovascular imaging specialist blinded to the treatment groups (Precession by Heart Imaging Technologies).

Hemodynamic Evaluation: Six weeks after PAB, pigs were anesthetized for right heart catheterization and venous blood collection. A Swan-Ganz catheter (Edwards Lifesciences) was advanced through into the pulmonary artery to measure RV pressure and cardiac output. After the procedure, animals were humanely euthanized for tissue collection. This study was approved by the University of Minnesota Institutional Animal Care and Use Committee (IACUC).

RA histological evaluation: Right atria specimens were fixed in 10% formalin, embedded in paraffin, and sectioned at 10 μm. Sections were subjected to deparaffinization and rehydration then stained with Alexa488 Wheat Germ Agglutinin and DAPI. Images were collected on a Zeiss LSM900 Airyscan 2 confocal microscope. Cardiomyocyte cross sectional area was blindly quantified by RTM using FIJI.

snRNAseq: RA nuclei were isolated as we have described using a protocol for frozen cardiac tissue(1,5). Nuclei were isolated from four PAB and four control pigs to prevent batch effects and sequence all nuclei at the same time. Sequencing and genome alignment (Sscrofa10.2) was performed by the University of Minnesota Genomics Center.

Analysis of snRNAseq data: snRNAseq analysis was performed using Minnesota Supercomputing Institute Rstudio v4.4 following our previously used Seurat workflow(1,6). After filtering, doublet removal, normalization, feature selection, scaling and dimensionality reduction, Azimuth(7-10) was used to guide clustering. We initially identified 26 clusters and determined cell types at the cluster level. Three clusters did not contain genes highly expressed in any cell type in the Single Cell Portal(11,12), were determined to be noise and removed from subsequent analysis. Differential gene expression analysis was performed using DESeq2(13) after nuclei were pseudobulked, and genes were considered differentially expressed if |log2FoldChange| ≥ 0.5 and adjusted *p*-value <0.05. Pathway analysis of differentially expressed genes was performed using ShinyGO(14). CellChat v2, following their standard workflow, evaluated predicted cellular communication(15).

Quantitative mitochondrial proteomics analysis: Mitochondrial enrichments from four control and four PAB RA and RVs were performed as previously described using the Mitochondrial Isolation Kit (Abcam). TMT16-plex proteomics (*n*=4 control RA, *n*=4 PAB RA, *n*=4 control RV, and *n*=4 PAB RV) was performed as described(16) to define the relative abundances of proteins in each sample.

Analysis of proteomic data: Relative abundances of proteins was determined using Proteome Discoverer 3.1 software. Then proteins that were significantly associated (*p*<0.05, Pearson test) with RA ejection fraction were subjected to KEGG pathway analysis using ShinyGO 0.82 online software (https://bioinformatics.sdstate.edu/go/).

Metabolomics analysis: Five control and five PAB RA frozen specimens were evaluated by Metabolon, Inc (Durham, NC) as reported(1,17,18).

Analysis of metabolomic data: Batch-normalized data were used for downstream analysis. Metabolites correlated to RA ejection fraction (|*r*| > 0.5) were selected for a pathway analysis using Metaboanalyst(19).

Western blot analysis: RA samples were extracted and 25 μg of total RA protein extract was subjected to SDS-PAGE. Western blots were analyzed with Odyssey Infrared Imaging system (Lincoln NE) was described(17). The Coomassie brilliant blue post transfer gel was scanned at the 700-nm wavelength and used as the loading control.

Electron microscopy: RA free wall tissue was placed into fixative (4% paraformaldehyde + 1% glutaraldehyde in 0.1M phosphate buffered, pH 7.2 (PB)). After fixation, tissue was washed with PB, stained with 1% osmium tetroxide, washed in H_2_O, stained in 2% uranyl acetate, washed in H_2_O, dehydrated through a graded series of ethanol and acetone and embedded in Embed 812 resin. Following a 24-hour polymerization at 60°C, 0.1 µM ultrathin sections were prepared and post-stained with lead citrate. Micrographs were acquired using a JEOL 1400 Plus transmission electron microscope (JEOL, Inc., Peabody, MA) at 80 kV equipped with a Gatan Orius camera (Gatan, Inc., Warrendale, PA) at the Mayo Clinic(16).

Evaluation of mitochondrial morphology in electron micrographs: Cardiomyocyte mitochondria were blindly analyzed by RMM. Briefly, mitochondrial cristae were graded based on the percentage of mitochondria occupied by cristae and their intactness as described(20).

iPSC-ACM differentiation and tunicamycin treatment: Human induced pluripotent stem cells (iPSC, WTC, Allen Institute) were differentiated into atrial cardiomyocytes using STEMdiff Atrial Cardiomyocyte Differentiation Kit (STEMCELL Technologies). Cells were seeded (470,000 cells/well) onto hESC-Qualified Matrigel (337 µg/mL, Corning) coated 12-well dish. At 70-85% confluency cells were treated with Medium A. 48 hours later cell media was replaced with Medium B. After 48 hours media was changed to Medium C. 48 hours later media was changed to Medium C. Media was subsequently changed every 48 hours with fresh Maintenance Medium. Differentiated iPSC-ACMs were passaged using the STEMdiff Dissociation Kit (STEMCELL Technologies). 24 hours after passaging, the cell media was replaced with Maintenance Media and treated with either tunicamycin (5 μg/mL, Sigma) or vehicle control (DMSO) overnight.

Mitochondrial network analysis: iPSC-ACM plated on chamberslides were stained with MitoTracker Orange (Thermoscientific) and superresolution confocal micrographs were captured on a Zeiss Airyscan 2.0 as described. Mitochondrial network morphology was blindly quantified using the Mitochondrial Network Analysis toolset(21) by MK.

Seahorse analysis: Following overnight tunicamycin treatment, mitochondrial respiration was assessed using a Seahorse XFe96/XF Pro Extracellular Flux Assay Kit (Agilent). 30 minutes prior to the assay, the cell culture medium was changed to XF RPMI Medium (Agilent) containing 5 mM glucose, 4 mM glutamine, and 1 mM pyruvate. The final concentrations of inhibitors were 5 uM/well oligomycin (Sigma), 20 uM/well FCCP (Cayman), and 1 μM/well antimycin/rotenone (Sigma). OCR measurements were normalized to total number of cells per well by measuring nuclear intensity. After the assay, cells were stained with Hoechst (1μg/mL, Biotium). Micrographs were taken on a Zeiss Axio Observer Z1 microscope and analyzed for fluorescence intensity using FIJI.

Data Availability: snRNAseq data are available on NCBI Gene Expression Omnibus. Proteomics data are available on Figshare.

Supplemental Figure 1: Immunoblots confirmed downregulation of proteins involved in endoplasmic reticulum homeostasis in PAB RA specimens. (A) Representative Western blots and subsequent quantification (B) of endoplasmic reticulum proteins in control and PAB right atrial samples. P-values determined by unpaired t-test.

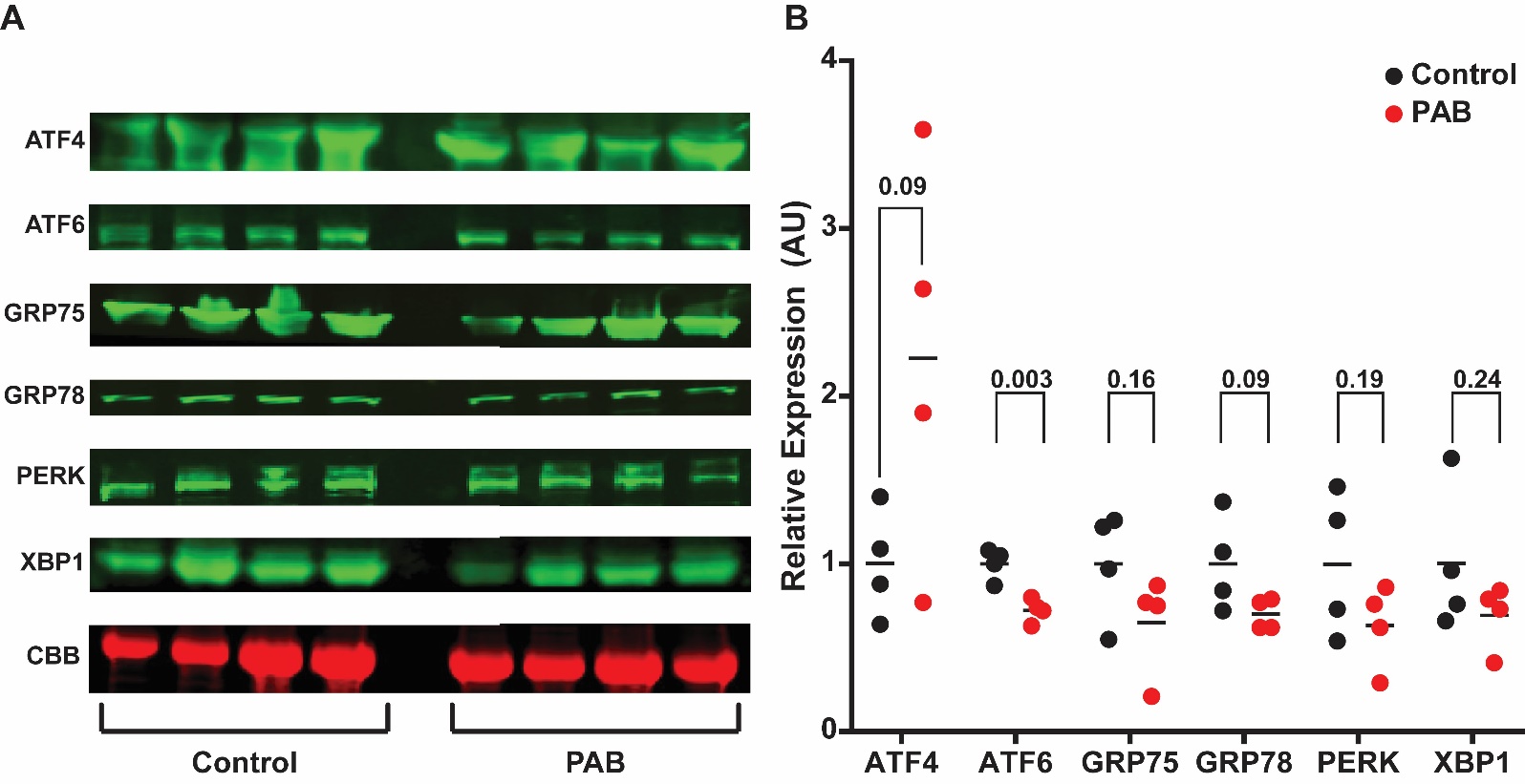

Supplemental Figure 2: snRNAseq revealed the dysfunctional RA had an accumulation of pro-inflammatory macrophage. (A) UMAP of total macrophage subcluster. (B) UMAP of control macrophage subclusters. (C) UMAP of PAB macrophage subcluster. (D) Quantification of relative abundance of three macrophage clusters identified. (E) Pathway analysis of transcripts upregulation in PAB macrophage. (F) Pathway analysis of downregulated transcripts in PAB macrophage. (G) Cell-chat analysis depicted the predicted number of cell-cell communications from macrophage to other cell types. (F) Predicted strength of interactions between macrophage and other cell types. Macrophage-cardiomyocyte interaction strength was predicted to be greater in PAB RA samples.

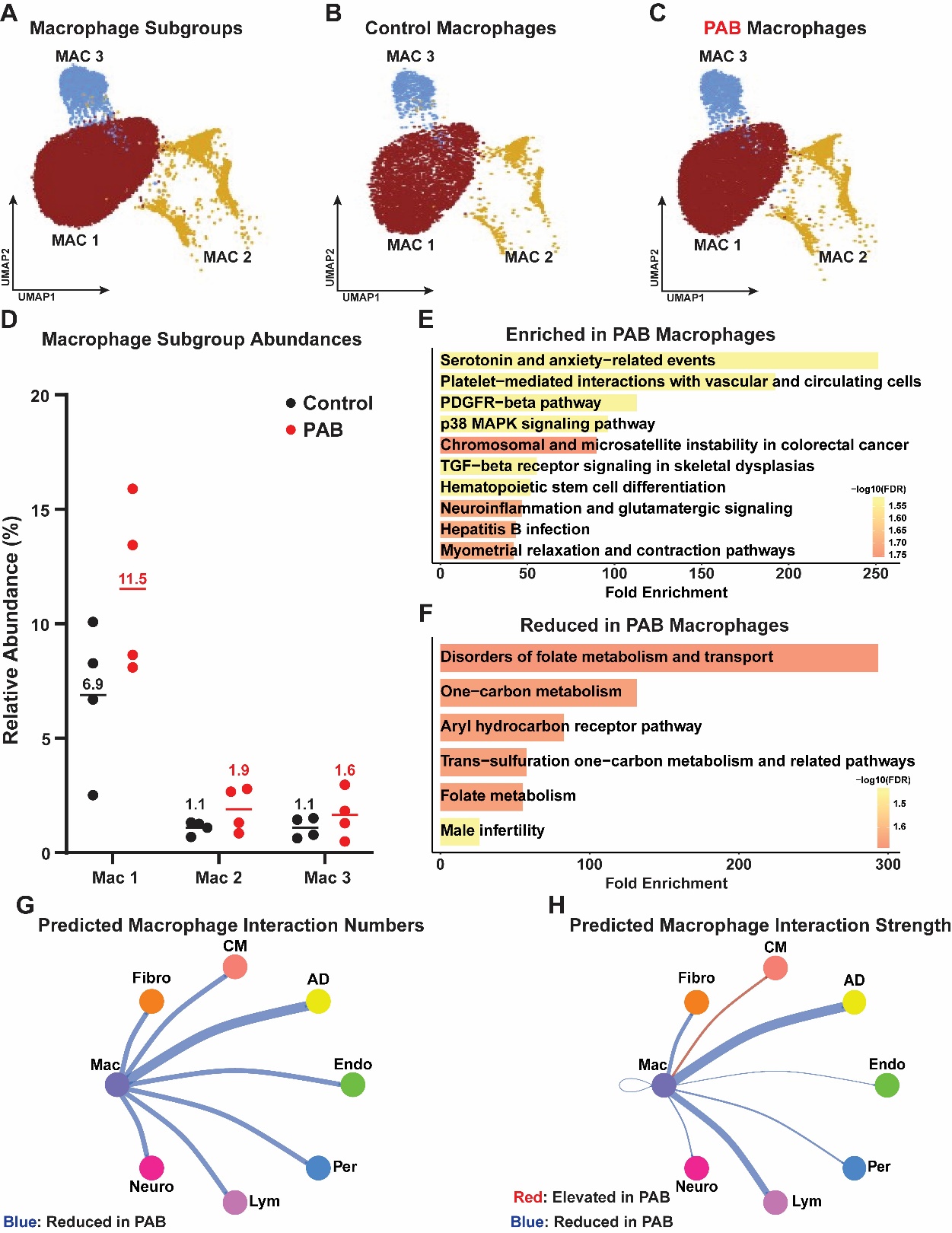

Supplemental Figure 3: PAB RA cardiomyocytes exhibited disruptions in mitochondrial cristae structure. (A) Representative electron micrograph of RA cardiomyocyte mitochondria. Red-arrows highlight mitochondria with disrupted cristae morphology. (B) Quantification of RA cristae score. *p*-value determined by Mann-Whitney test.

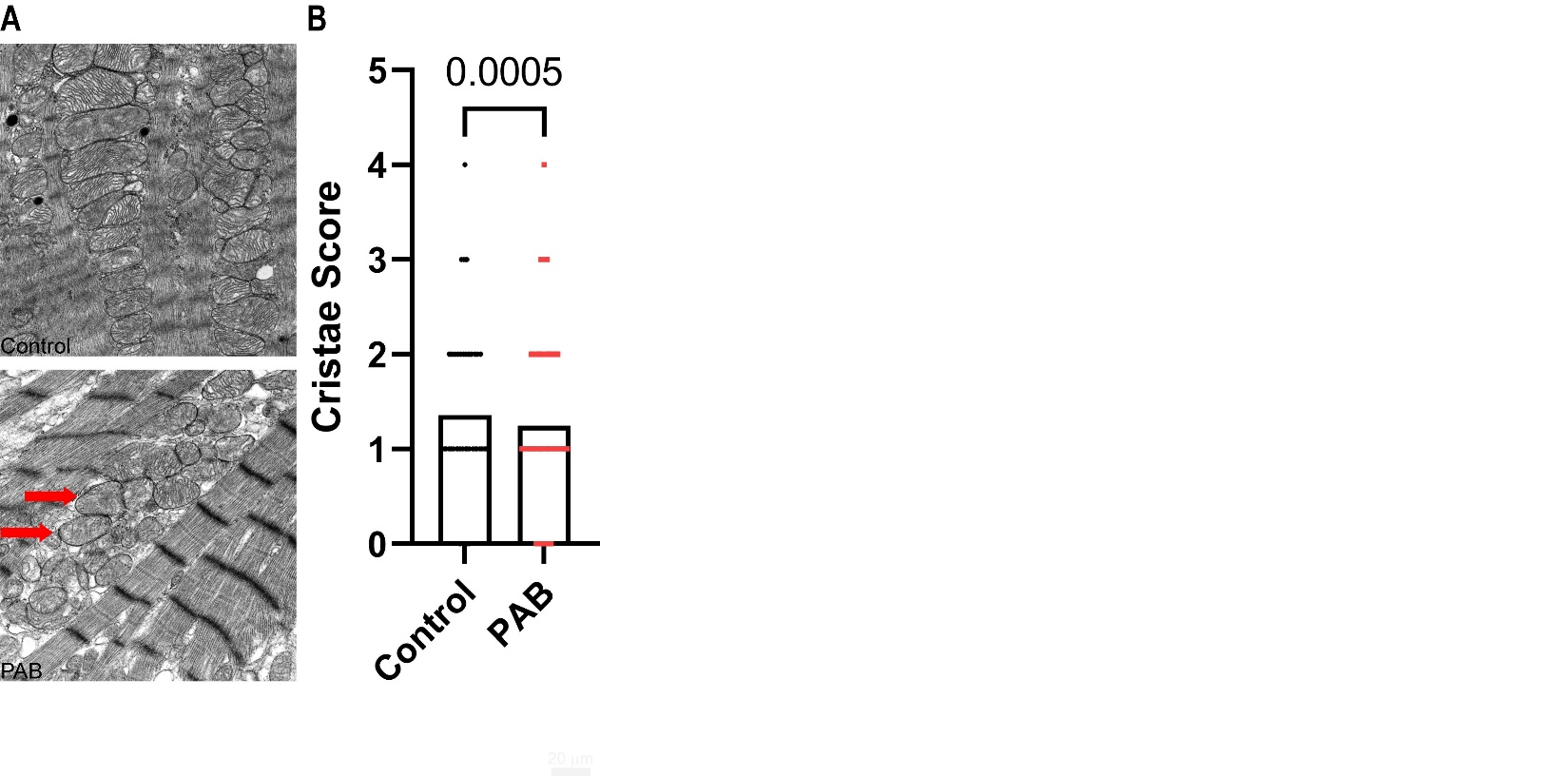

Supplemental Figure 4: Induction of ER stress with tunicamycin disrupted mitochondrial morphology and impaired mitochondrial oxidative capacity in iPSC-ACMs. (A) Representative image of iPSC-ACMs stained with the atrial-specific marker (MLC-2A, Synaptic Systems, demonstrating differentiation into atrial cells. (B) Representative confocal micrographs of iPSC-ACMs stained with MitoTracker Orange. Arrows indicate mitochondrial doughnuts Tunicamycin induced mitochondrial dysregulation as evidenced by a reduction in mitochondrial footprint (C) and an increase in mitochondrial doughnuts (D). *p*-values determined by Mann-Whitney test. (E) Representative Seahorse tracings of oxygen consumption rates. Tunicamycin impaired mitochondrial function as demonstrated by a significant reduction in basal oxygen consumption rates (F), maximal oxygen consumption rates (G) and ATP-linked oxygen consumption rates (H). *p*-values determined by unpaired t-test or Mann-Whitney test.

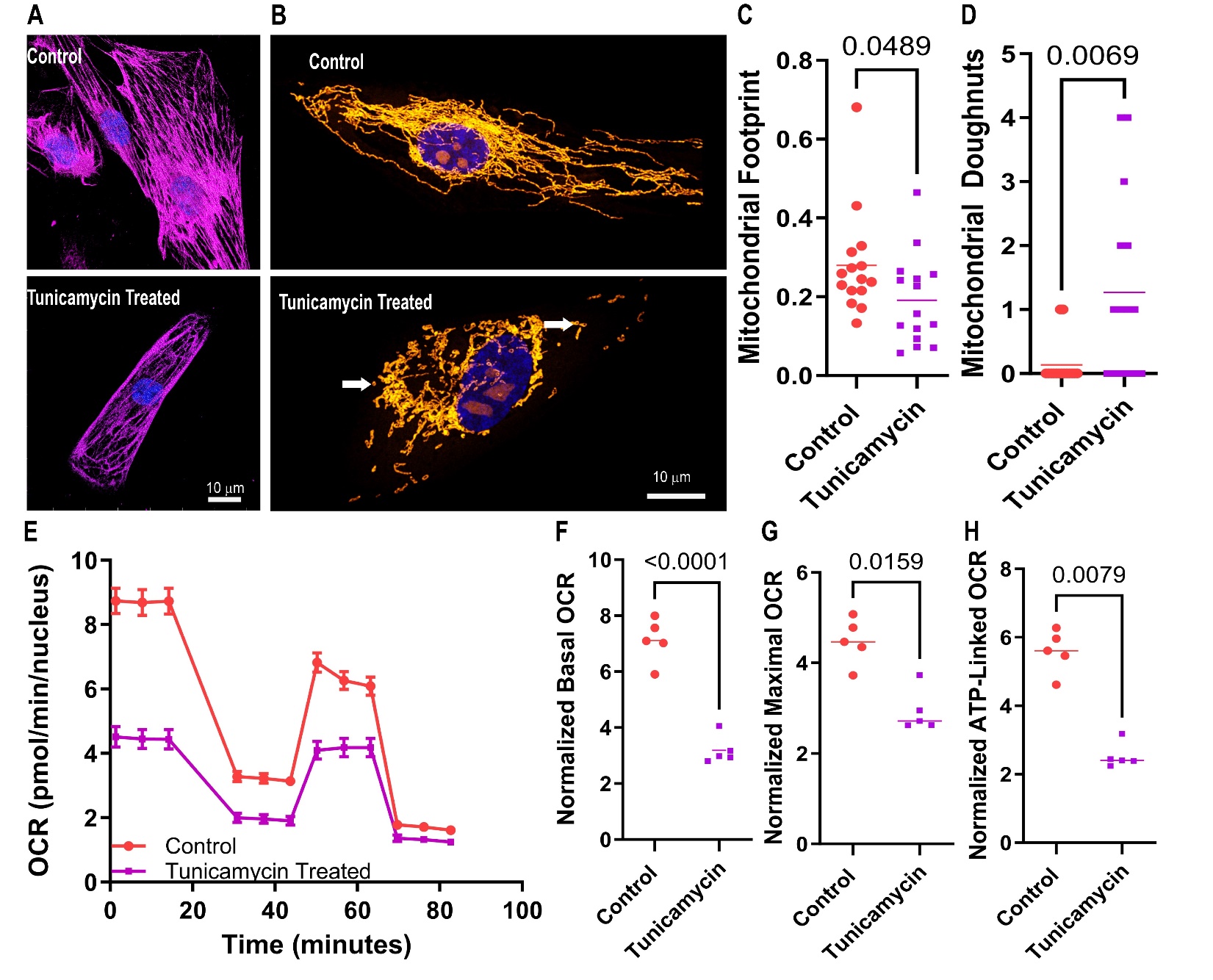

Supplemental Figure 5: Examination of BCAA catabolic enzyme transcripts in the RA and RV from the human heart atlas (A) and BCAA metabolism enzyme relative abundances in porcine proteomics experiments in the RA and RV (B) using Volcano plots. No proteins were significantly different using a FDR of 0.05 (dotted line). BCAA degrading enzymes were not expressed at higher levels in the RA compared to the RV in either human or porcine hearts.

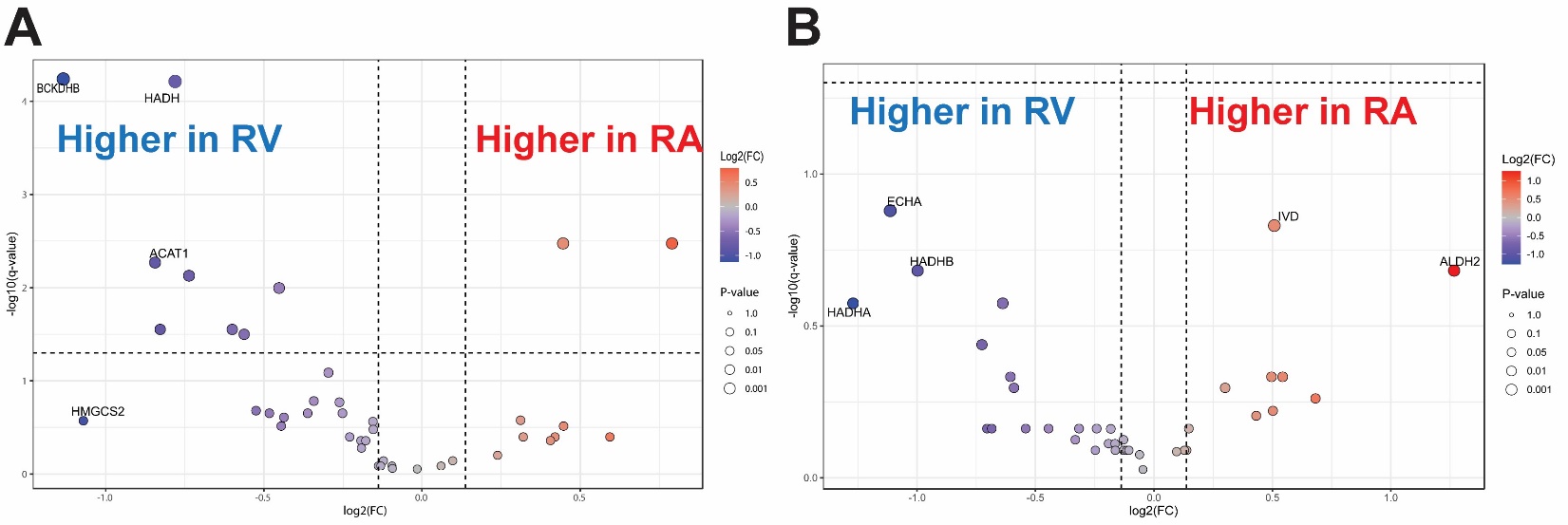

Supplemental Figure 6: BMP10 regulation was not altered in PAB RA samples. (A) UMAP with relative levels of BMP10 transcripts in all cell types. BMP were predominately detected in cardiomyocyte nuclei. (B) Quantification of BMP10 expression in cardiomyocytes. (C) UMAP depicting BMP10 expression in control cardiomyocytes. (D) UMAP depicting BMP10 expression in PAB cardiomyocytes.

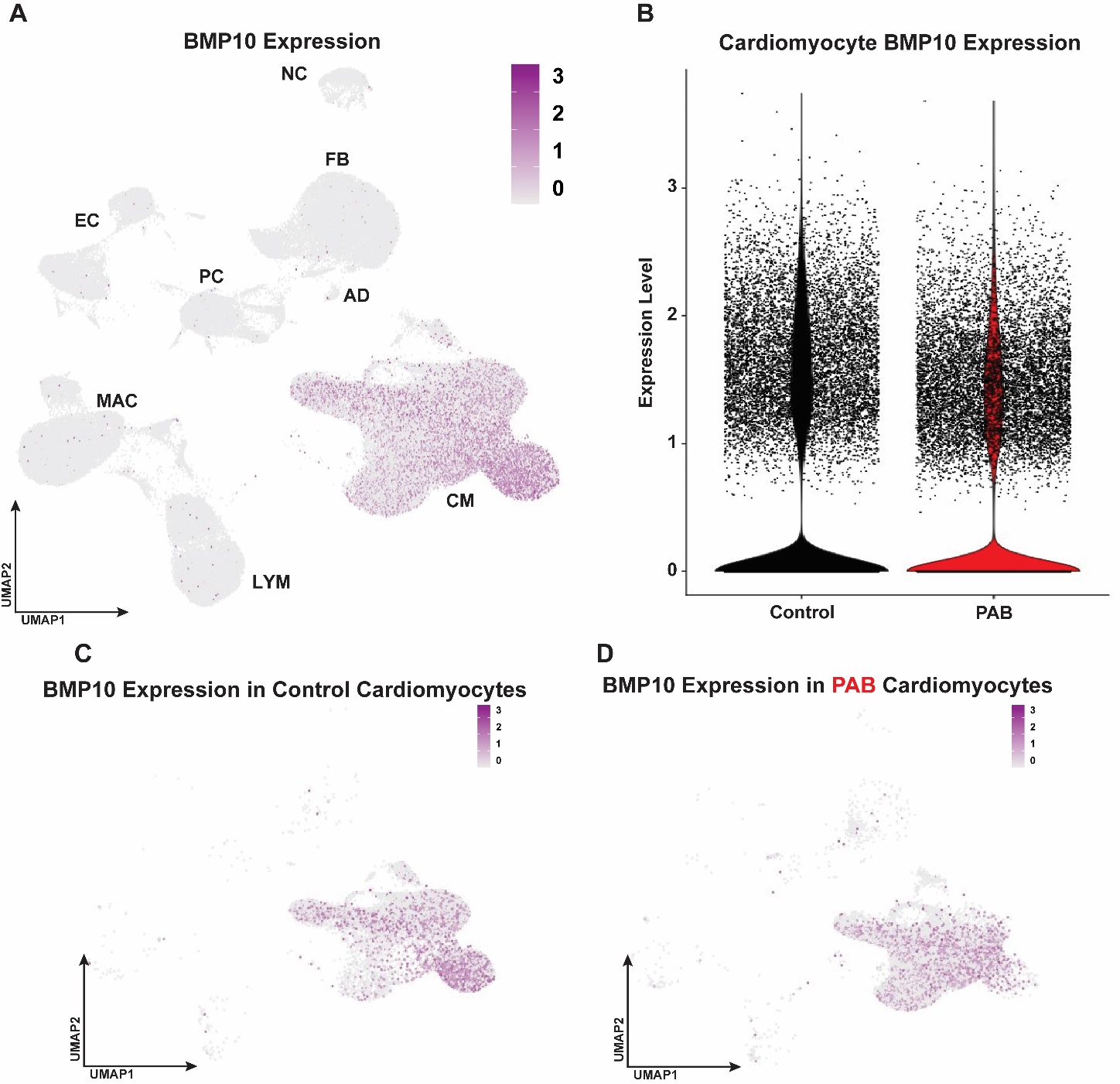

Supplemental Table 1: List of antibodies used

| Antigen | Company | Catalogue Number | Dilution | Experimental Approach |
| --- | --- | --- | --- | --- |
| ATF4 | Proteintech | 10835-1-AP | 1:250 | Western blot |
| ATF6 | Proteintech | 24169-1-AP | 1:250 | Western blot |
| GRP75 | Cell Signaling | 3593T | 1:250 | Western blot |
| GRP78 | Abcam | Ab21685 | 1:250 | Western blot |
| PERK | Cell Signaling | 3192T | 1:250 | Western blot |
| XBP-1s | Cell Signaling | 12782T | 1:200 | Western blot |
| MLC | Synaptic Systems | 311 011 | 1:250 | Immunofluorescence |
| Anti-rabbit Alexa 555 secondary antibody | Invitrogen | A32732 | 1:500 | Immunofluorescence |
| Anti-mouse secondary | Li-COR | 926-32210  926-68070 | 1:5000 | Western blot |
| Anti-rabbit secondary | Li-COR | 926-32211  926-68071 | 1:5000 | Western blot |
